# Phenology of questing *Ixodes frontalis* nymphs reveals a late-winter to early-spring activity peak

**DOI:** 10.64898/2025.12.23.696149

**Authors:** Olivier Plantard, Camille V. Migné, Sara Moutailler, Frédéric Stachurski, Albert Agoulon

## Abstract

Although *Ixodes frontalis* is widespread across Europe, including urban and peri-urban habitats, several aspects of its biology remain undescribed. One such aspect is the population dynamics of its nymphal stage, which has so far been reported from limited sample sizes, preventing the identification of its peak abundance. To fill this knowledge gap, we conducted a year-long monthly survey at four locations in France representing three distinct climates. The sampling protocol—based on the flagging method performed under bamboo bushes where larvae of this species had previously been found—was designed to maximize the likelihood of capturing the nymphal stage. Based on the 400 nymphs collected during this survey, a clear peak of *I. frontalis* nymphs was observed around March (February to April) across all four sites, with an additional peak in August at one site. The population dynamics of the larval stage—based on semi-quantitative counts—showed the same previously described pattern, with a peak in late autumn. Twenty adults (11 females and 9 males) were also collected from March to December, but these data did not allow a clear identification of a peak abundance. Because several bird species—such as blackbirds, song thrushes, European robins, and Eurasian blackcaps—serve as feeding hosts for both *I. ricinus* and *I. frontalis*, and are also reservoirs for pathogens transmitted by these ticks, the implications of our findings for the eco-epidemiology of tick-borne diseases are discussed.

**Graphical abstract:** 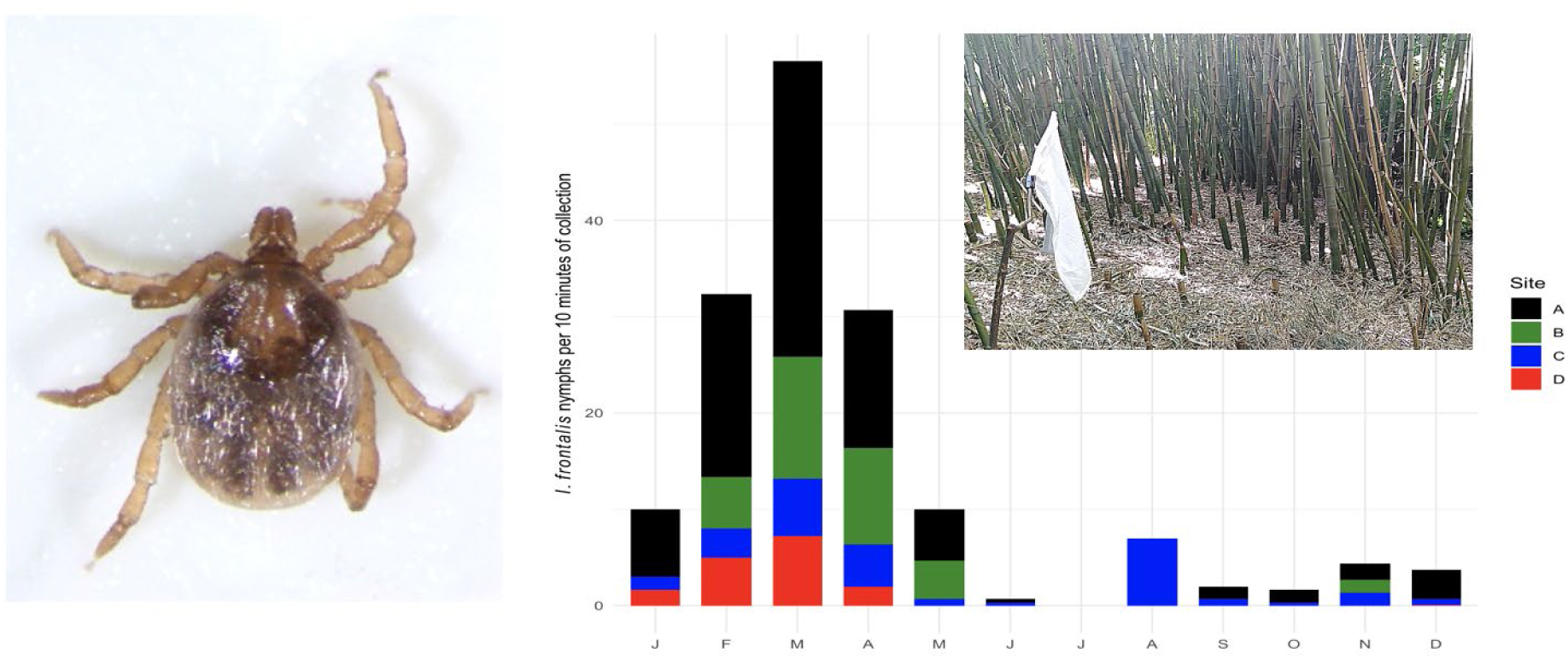

**Highlights:** A sampling method for *Ixodes frontalis* nymphs was developed

Four hundred nymphs were collected across four different sites in France through a monthly survey

A clear peak was observed around March, with an additional peak in August at one site

## Introduction

Besides small mammals and ruminants, passerine birds are considered important hosts for numerous tick species, both as feeding and reservoir hosts, but also due to their high dispersal ability (Norte et al. 2013, Heylen 2016, Rataud et al. 2024). This is particularly true for *Ixodes ricinus*, the most abundant tick species in Europe and the main vector of *Borrelia burgdorferi* sensu lato, the agent of Lyme disease in humans (Rataud et al. 2022). Among the different *Borrelia* species known to cause Lyme disease in humans, *B. garinii* and *B. valaisiana* are considered to have their main reservoirs in birds, which also harbor other *Borrelia* species with unknown human pathogenicity, such as *B. turdi* (Coipan & Sprong 2016, Eisen 2020). However, birds are also commonly parasitized by other tick species, notably by *I. frontalis*, an ornithophilic tick. Unlike *I. ricinus*, experiments on the vector competence of *I. frontalis* have shown that it is not able to transmit *B. garinii* (Heylen et al. 2014). *Ixodes frontalis* is also known to harbour *Anaplasma phagocytophilum* (Jahfari et al. 2014, Wilhelmsson et al. 2024). Although it is widespread throughout the Western Palearctic region, the biology of *I. frontalis*, and particularly its phenology, remains poorly known, except for the larval stage (Agoulon et al. 2019). Understanding the phenology of a given tick species is essential to assess its role in the eco-epidemiology of tick-borne diseases, as it determines which host species are available for blood feeding, particularly in ornithophilic species, where many migratory birds may be present or absent depending on the timing of each tick stage. Moreover, knowing the timing of peak abundance enables more efficient sampling, which is crucial for studies requiring large numbers of ticks, such as investigations on pathogen prevalence. Most available data on the phenology of *I. frontalis* come from collections of ticks found on birds. Such collection campaigns are not conducted evenly across the four seasons or among all bird species; they occur more frequently during the pre- and post-nuptial periods on species belonging to the *Passeriformes* order (especially migratory species). For this reason, the phenology inferred from such datasets may be biased. Finally, the number of *I. frontalis* collected during bird-capture-based surveys remains limited, preventing the clear identification of the timing of its abundance peak. With at least 73 bird species reported as hosts for *I. frontalis*, including 27 for nymphs (Keve et al. 2024), the sample size per bird species remains limited.

The population dynamics of *I. frontalis* larvae was first described through monthly tick collections at a site in Western France dedicated to the study of *I. ricinus* temporal dynamics (Wongnak et al. 2022). At this site, *I. frontalis* was incidentally discovered due to the presence of bamboo groves. In this particular habitat, *I. frontalis* larvae showed a peak in abundance in October–November, followed by a gradual decline during the subsequent winter months (Agoulon et al. 2019). Following this discovery, a second study revealed the presence of *I. frontalis* at nearly all sites containing bamboo groves (29 out of 32 sites surveyed) through tick collections conducted during its larval season (Plantard et al. 2021). However, only a small number of nymphs were collected in this latter study, conducted only between November and April. In the present study, a new sampling protocol was implemented with the primary objective of maximizing the collection of *I. frontalis* nymphs to determine their temporal dynamics. The secondary objective was to validate the larval dynamics observed in the first study (Agoulon et al. 2019) at new sites and in a different year.

## Material and methods

### Study sites

Among the 29 sites where *I. frontalis* larvae were previously found under bamboo groves (Plantard et al., 2021), three were selected for the present study due to their high number of nymphal stages (two sites near Nantes: sites A and B, and one near Paris: site C). A fourth site, located near Montpellier (site D), was added to include a Mediterranean climate (Table 1), as different climates may induce variations in phenology within a given tick species (see for example Hoch et al. 2024 for the case of *I. ricinus* or Allen 2025 for *I. scapularis*). Those four sites therefore encompass different climatic conditions: oceanic (Nantes), degraded oceanic (Paris) and Mediterranean (Montpellier) (Figure 1). At each of the four sites, ticks were collected monthly for at least one year (Table 1).

**Figure 1:**
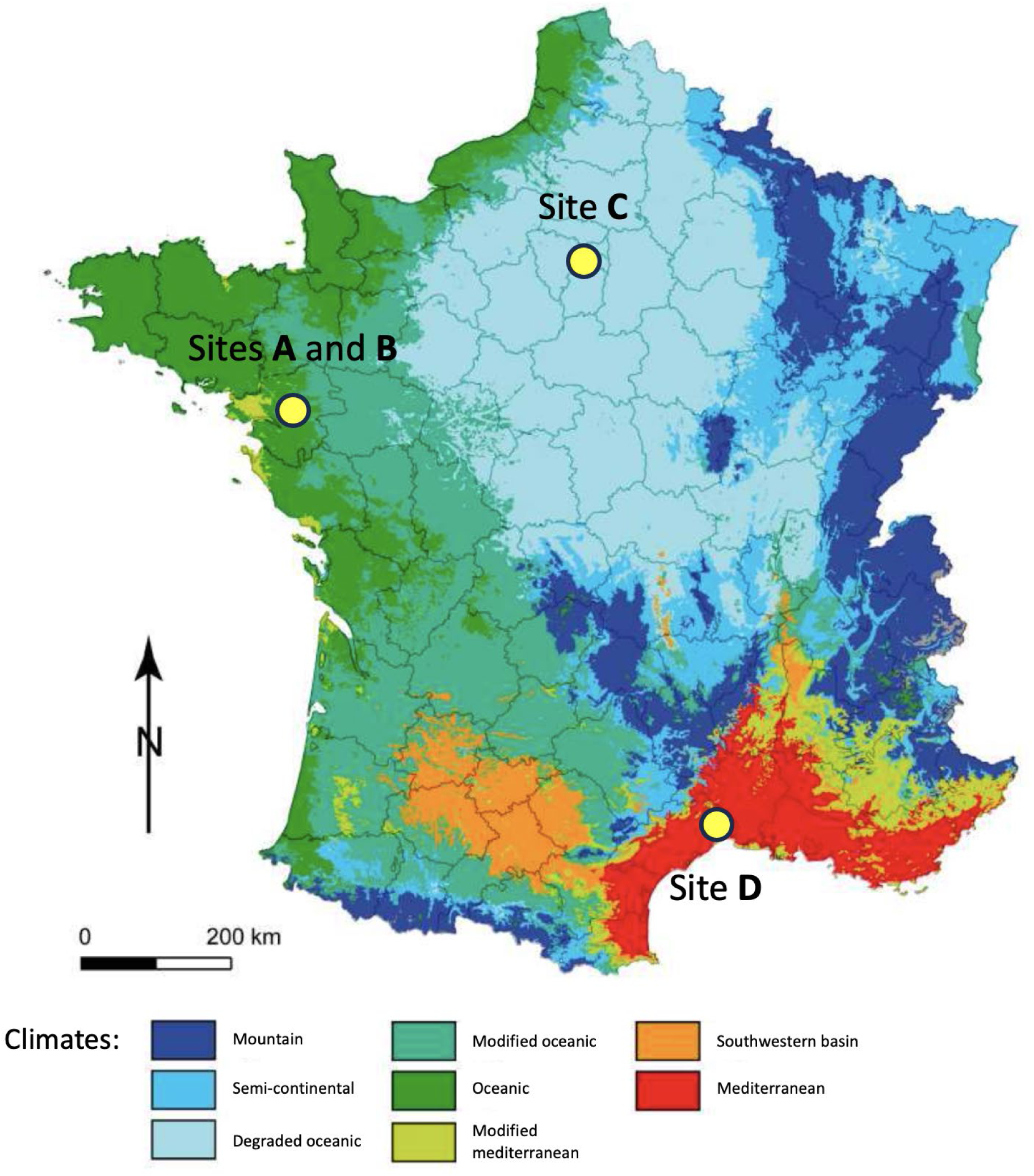
Localisation of the four study sites on a climate map of France (from Joly et al. 2010) Sites A (47°14’37"N, 1°36’59"W) and B (47°17’18"N, 1°31’31"W) are near Nantes Site C (48°50’15"N, 2°26’37"E) is near Paris Site D (43°41’29"N, 3°52’39"E) is near Montpellier

**Table 1.**
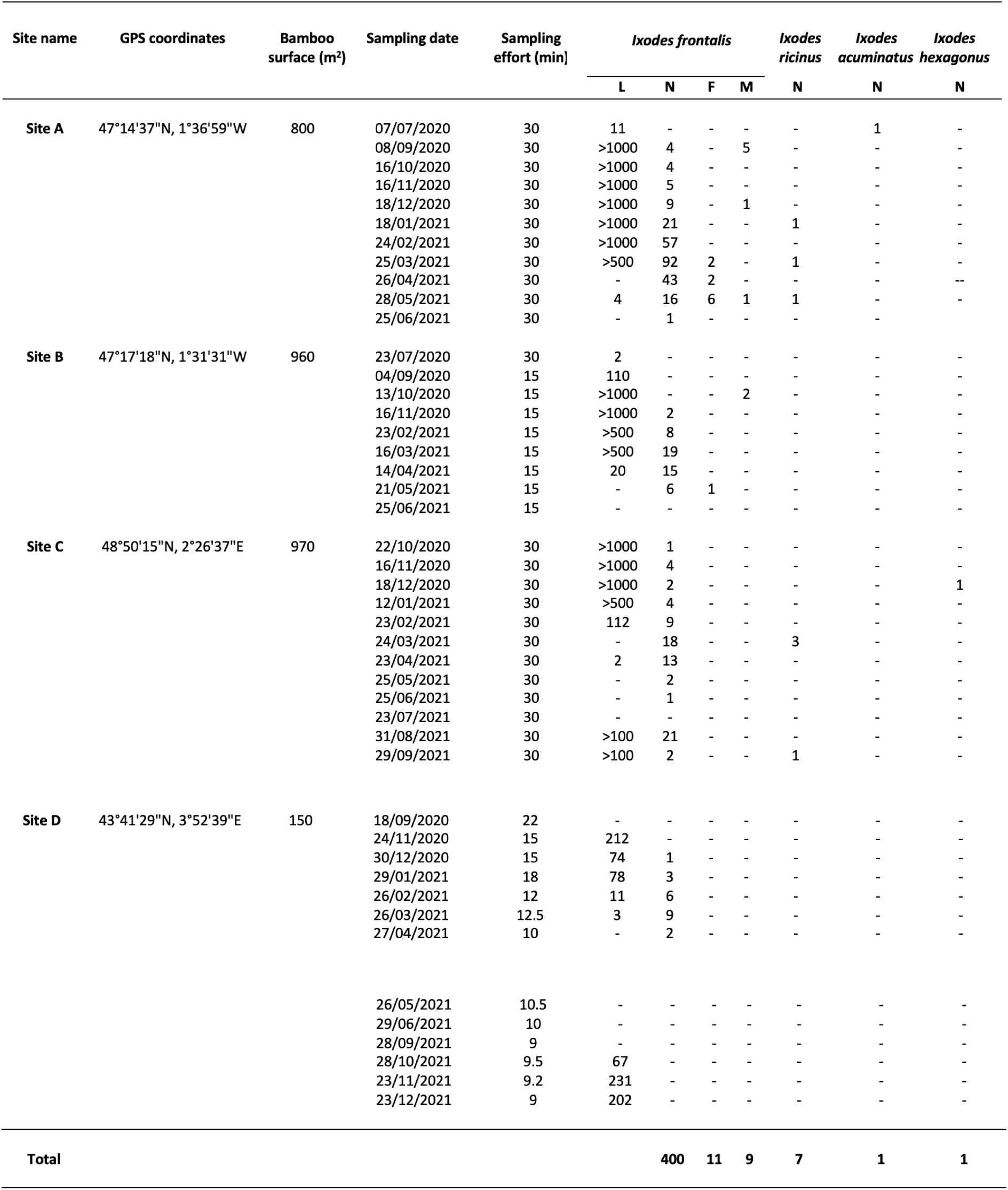
Collection data of questing ticks across the 4 study sites (L=larva, N=nymph, F=female, M=male, for each tick species).

### Tick sampling and identification

Because the dragging method is nearly impossible to perform in bamboo groves—where bamboo stalks are typically spaced less than one meter apart (see supplementary information in Plantard et al. 2021), i.e., the standard width of a drag sheet—the flagging technique was used as described by Plantard et al. (2021), with a modification in how sampling time was measured. Specifically, to estimate the relative density of nymphs, only the duration of active collection was considered. Using a stopwatch, time was recorded solely when the flag was in contact with the ground. During visual inspection of the flag (i.e., when it was held at eye level) or during tick removal, the stopwatch was paused. Ticks were removed from the flag using tweezers and placed in Eppendorf tubes for subsequent identification in the laboratory. To standardize collection effort, the number of nymphs was expressed per 10 minutes of collection time. Because flagging efficiency may vary between collectors, the same person performed all monthly surveys at each site. Analyses therefore focused on monthly variations in tick density within a given site, without considering differences in tick density among sites. As the surface area of the bamboo groves differed between sites—ranging from 150 to 800 m²—the total collection time per site also varied (Table 1). All nymphs and adults were identified in the laboratory under a stereomicroscope, using the identification keys from Estrada-Peña et al. (2017) and Pérez-Eid (2007).

### Assessment of larval abundance

Although the present study focused on the nymphal stage, we took advantage of the monthly survey at sites where *I. frontalis* was present to confirm, using this new protocol based on the flagging technique (rather than the blanket-dragging method described by Agoulon et al. 2019), the phenology of the larval stage at four additional sites, including two distant ones representing new climatic conditions (sites C and D). In the present study, the large numbers of larvae observed during the autumn and winter months (sometimes several thousand individuals) did not always allow for exhaustive counting or collection of this stage. Because precise counts were not required to validate the general population dynamics of *I. frontalis* larvae, a semi-quantitative assessment was used when larvae were present in large numbers (>100, >500, >1000). To verify that these numerous larvae belonged to *I. frontalis*, a subsample of 100 larvae from each monthly sample—when full collection and identification were not possible—was placed in Eppendorf tubes containing 70 % ethanol for species confirmation by morphological examination under a microscope, using the same identification keys as above.

## Results

### Diversity of tick species

During the study, four tick species were found beneath the bamboo bushes. *Ixodes frontalis* was by far the most abundant (11 females, 9 males, 400 nymphs, and >14 000 larvae) and was present at all four sites. It was followed by *I. ricinus* (7 nymphs) found at sites A (n = 3) and C (n = 4), *I. acuminatus* (1 nymph) found at site A, and *I. hexagonus* (1 nymph) found at site C (Table 1).

### *Distribution of* Ixodes frontalis *life stages across sites*

The distribution of *I. frontalis* across sites was as follows: at site A, 10 females, 7 males, 252 nymphs, and >6 500 larvae were recorded (identification confirmed for 100 % of a subsample of 715 larvae); at site B, 1 female, 2 males, 50 nymphs, and >3 100 larvae (confirmation rate 100 % for 532 larvae); at site C, 77 nymphs and >3 800 larvae (confirmation rate 100 % for 714 larvae); and at site D, 21 nymphs and 878 larvae (all individuals counted and identified).

### *Description of* Ixodes frontalis *nymphal phenology*

At all four investigated sites, *I. frontalis* nymphs displayed a pronounced peak in March, with the highest densities observed from February to April, and an additional late-summer peak in August observed only at site C (Figure 2). After standardizing the raw nymph counts by collection time to estimate the number of nymphs per 10 minutes of sampling, the March peak remained clearly dominant, followed by a secondary peak in August at site C (Figure 3). No nymphs were collected in July at any site, even at the most productive one (site A: total = 252 nymphs, with 92 collected in March 2021, corresponding to 30 nymphs per 10 minutes of sampling).

**Figure 2:**
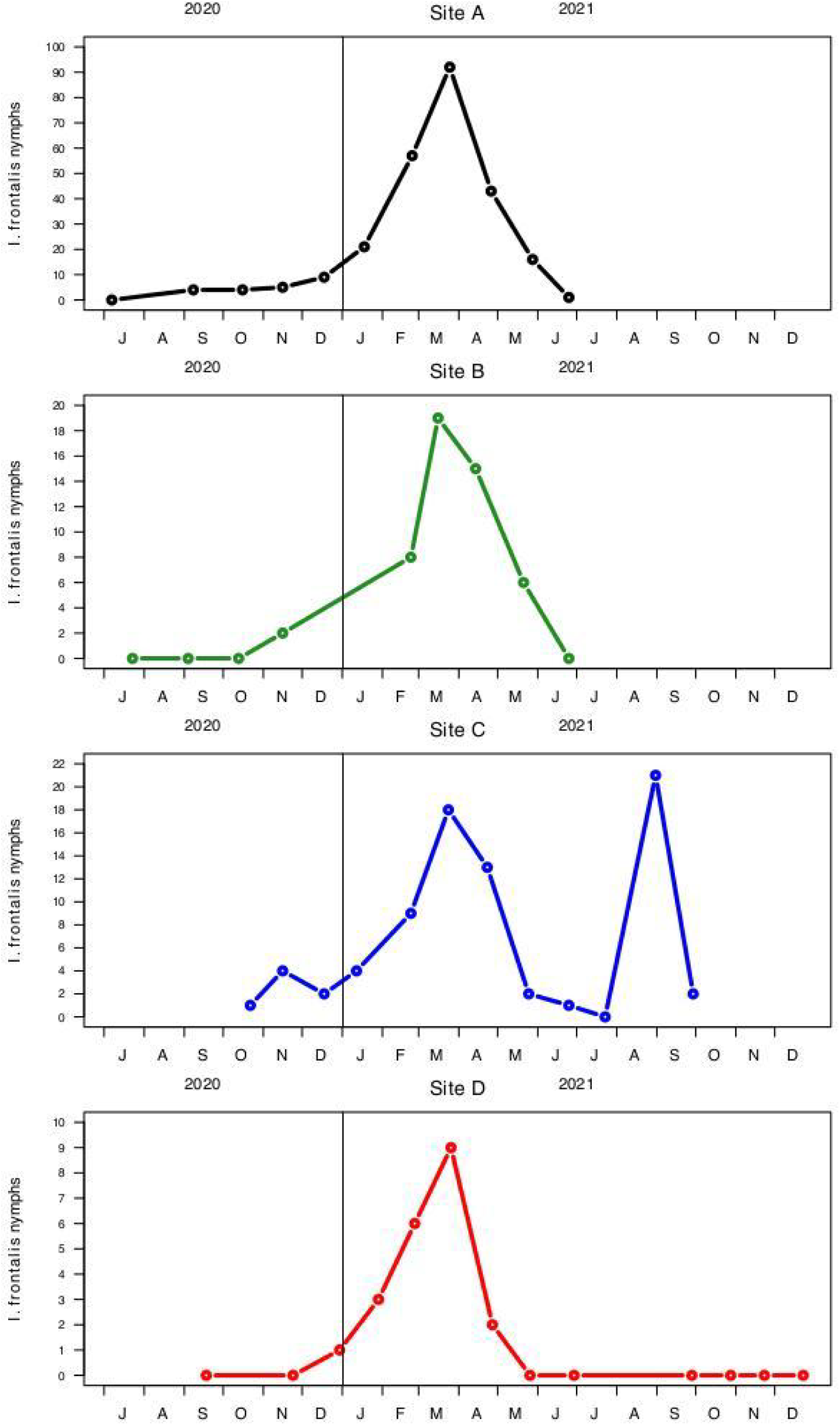
Monthly abundance of *Ixodes frontalis* nymphs on each study site.

**Figure 3:**
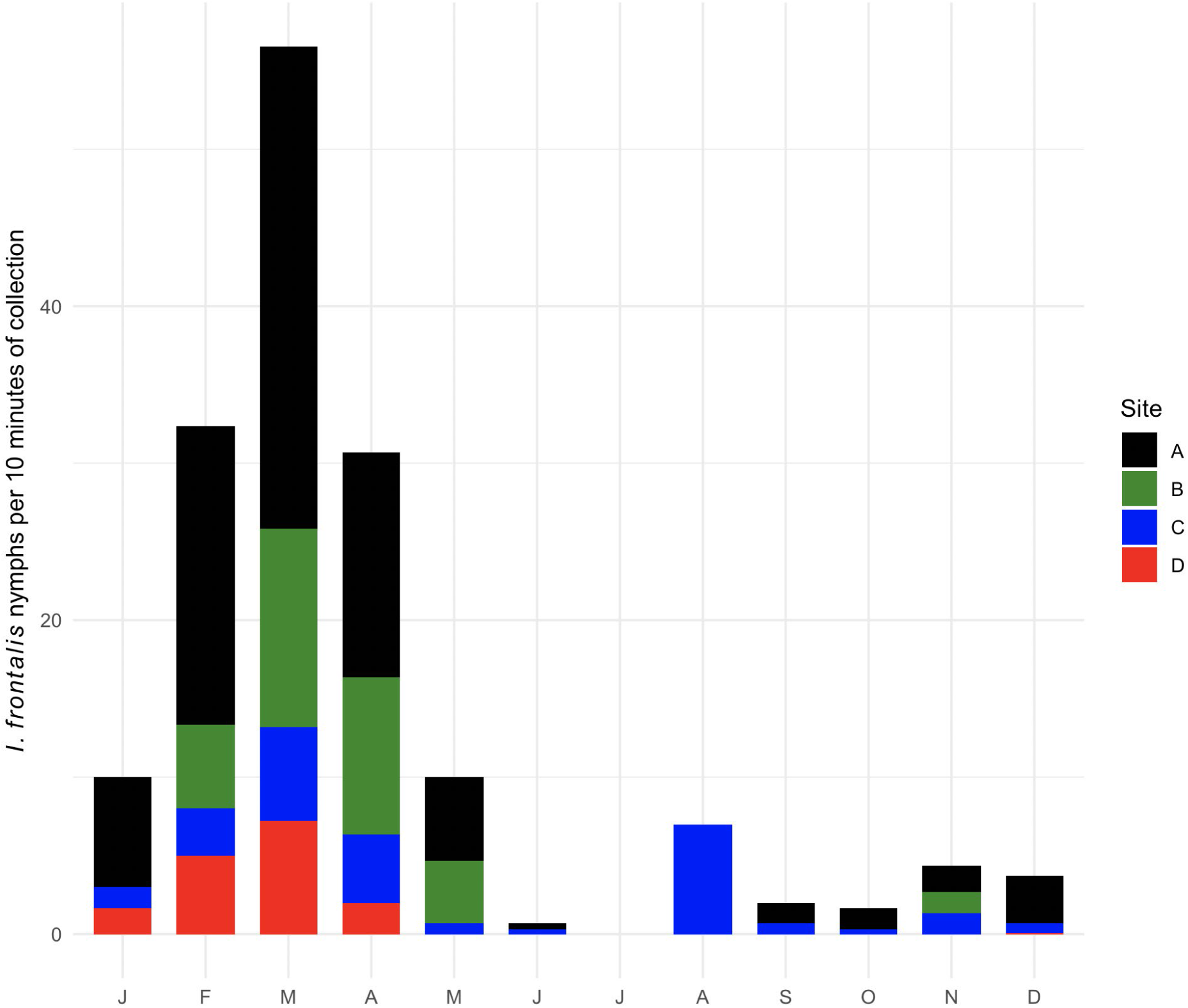
Standardized monthly abundance of *Ixodes frontalis* nymphs. For each month of the year (2020 and 2021 were pooled) and for each study site, the total number of collected nymphs was divided by the collection time in minutes and multiplied by 10, representing the number of nymphs per 10 minutes of collection.

### Description of Ixodes frontalis larval phenology

Larval sampling beneath the bamboo bushes revealed the exclusive presence of *I. frontalis* (100 % of the 2 839 identified larvae). Larvae were present during the same period at all four sites, i.e., throughout the autumn–winter season, mainly between October and February. Larval numbers were estimated to exceed one thousand during at least two months (site B) and up to six months (site A) of this period, except at site D, where the maximum number of larvae reached only 212 individuals. Larval counts were zero or limited to only a few individuals between April and July at all sites (during this four-month period: total = 15, max = 11 at site A; total = 22, max = 20 at site B; and 2 at site C) (Table 1). The increase in larval numbers observed between late summer and early autumn was steep, suggesting a sudden onset, whereas the decline between late winter and early spring was more gradual, indicating a progressive disappearance of larvae over time.

## Discussion

### Diversity of tick species

As previously documented (Agoulon et al. 2019; Plantard et al. 2021), the bamboo bush habitat appears to spatially concentrate *I. frontalis*, with thousands of larvae, hundreds of nymphs, and dozens of adults collected at the four study sites. However, this species is not strictly confined to this habitat and can also be found, for example, in large deciduous forests distant from urban areas, where its presence becomes evident when numerous larvae are collected on the flag or drag cloth during winter months (Plantard, pers. obs.). For nymphs and particularly adults, the flagging method yielded more individuals than the dragging technique (used in Agoulon et al. 2019), likely because the heavier stages—especially adults—tend to fall from the drag cloth more easily during 10 m drags, as previously reported for other tick species (Tack et al. 2011).

In these particular bamboo bush habitats, especially those located in urban areas, *I. ricinus* appears to be rare. Nonetheless, a few nymphs were recorded at sites A and C (site D being located in a Mediterranean area where *I. ricinus* is absent). Although not all tick larvae observed on the flag were collected and identified due to their large numbers, no *I. ricinus* larvae were identified among the 100 larvae examined each month during periods of high larval abundance. As the larval peak for *I. ricinus* occurs around June (Agoulon et al. 2019), the presence of this species’ larvae would have been easily detectable on the flag if they had been present. However, no larvae and only a very small number of nymphs (one each at sites A and C) were observed during this period. Because the four study sites were located near urban areas, the presence of roe deer—major hosts for *I. ricinus* females—is unlikely. Consequently, the probability that an engorged *I. ricinus* female would drop off within a bamboo grove is very low. Moreover, we hypothesize that rodent density is low in bamboo groves because this habitat has very sparse understory vegetation (see supplementary information, Plantard et al. 2021, site A). The soil is almost entirely covered by dead bamboo leaves, offering little food or shelter for small mammals. For these reasons, we hypothesize that the few *I. ricinus* nymphs collected originated from larvae that had fed on birds using the bamboo groves as roosting sites during autumn and winter, as previously suggested by Agoulon et al. (2019) and Plantard et al. (2021), and that subsequently dropped to the ground. Birds—especially passerines such as the common blackbird (*Turdus merula*), song thrush (*Turdus philomelos*), and European robin (*Erithacus rubecula*)—which frequently occur in these bamboo groves, have already been reported as hosts for *I. ricinus* larvae (e.g., Marsot et al. 2012; Rataud et al. 2022; Keve et al. 2022).

Although based on a single observation, the presence of an *I. acuminatus* nymph at site A is noteworthy. This species remains poorly known and is only rarely collected from vegetation using flagging or dragging methods (Gilot et al. 1992; but see Petney et al. 2015). Most available data come from host-based sampling, mainly on small rodents, as this species is considered endophilic and largely restricted to burrows (Pfäffle et al., in Estrada-Peña et al. 2017). However, a few records of *I. acuminatus* on birds—particularly passerines—have been reported (reviewed in Keve et al. 2022), mainly involving larvae found on ground-foraging species such as the Eurasian wren (*Troglodytes troglodytes*) and the common chaffinch (*Fringilla coelebs*). As for the few *I. ricinus* nymphs observed in the present study, we hypothesize that this *I. acuminatus* nymph originated from a larva that had fed on a bird and dropped from its host during a nocturnal stay within the bamboo grove.

For both *I. ricinus* and *I. acuminatus*, this hypothesis could be tested using molecular methods to identify the host of the last blood meal (Goethert et al. 2021). The presence of an *I. hexagonus* nymph at site C may reflect the hedgehog’s role in maintaining tick populations in urban areas—including both *I. hexagonus* and *I. ricinus*—in the absence of alternative hosts (Földvári et al. 2011).

The occurrence of several tick species in syntopy within bamboo groves could contribute to the maintenance and circulation of potential tick-borne pathogens in urban areas, with some species possibly acting as bridge vectors (Heylen et al. 2017). As at least three of the four tick species are suspected to feed on the same bird species that frequent bamboo groves, co-feeding might represent a relevant mechanism promoting pathogen spillover between tick species. However, for co-feeding to occur, the phenological stages involved must overlap in their periods of activity. This appears to be the case for *I. frontalis* larvae or nymphs and *I. ricinus* nymphs, for example in March.

### Ixodes frontalis *nymphal phenology*

Our previous investigation of *I. frontalis* phenology was based on a monthly survey using the dragging method applied to ten 10-m-long transects (i.e., 100 m²) with three successive passes (Agoulon et al. 2019). Although the species was abundant at that site—with several thousand larvae collected—only 28 nymphs of *I. frontalis* were recorded over the three years of sampling, spread across 17 months out of 36, with maxima of only three individuals (January or November 2015) or two individuals (March–April 2017). Those limited numbers were insufficient to determine the timing of the *I. frontalis* nymphal peak.

In the present study, the implementation of a new sampling design based on the flagging method in sites with a high density of *I. frontalis* larvae (i.e., bamboo groves) enabled the collection of a large number of nymphs and thus allowed clear identification of a nymphal peak in late winter to early spring (February – April), with the highest numbers recorded in March across all four investigated sites. The lowest abundances were observed in July (no nymphs at any of the four sites) and in June (a single nymph each at sites A and C). This timing of the nymphal peak agrees with the phenology described by Agoulon et al. (2019) and Remesar et al. (2023) (15 nymphs in November and 7 in February), or by Reynolds et al. (2022) (3 and 4 nymphs in March 2019 and 2020, 2 in September, 10 in February, 1 in April). However, the total number of *I. frontalis* nymphs collected was much higher in the present study (n = 400) than in the three previous ones (n = 28 over 36 months, n = 33 over 12 months, and n = 20 over 28 months, respectively). Fourteen and fifteen *I. frontalis* nymphs were recorded during two-year monthly surveys by Remesar (2019a, 2019b), although their timing was not reported. The occurrence of an additional peak in late August at one of our study sites (and the two September nymphs reported by Reynolds et al. 2022) suggests that a late-summer “secondary peak” may occur sporadically. The extended period during which nymphs were collected could be explained by (i) variability in the development time of engorged larvae, or (ii) the occurrence of diapause—either behavioral diapause in nymphs (periods of inactivity during the hottest months) or developmental diapause in larvae, resulting in delayed molting to the nymphal stage. Given the small size of the bamboo groves studied and the highly mobile nature of birds, a single bird landing at a site infested with engorged *I. frontalis* larvae—including migratory birds carrying ticks from distant regions with distinct phenologies—could account for the secondary peak observed in August. *Ixodes frontalis* is distributed from western Europe to central Asia, a vast range where intraspecific variability in phenology may occur. Furthermore, two genetically distinct mitochondrial haplotypes have been described within this species (Hornok et al. 2016; Charrier et al. 2019; Reynolds et al. 2022). It would therefore be of interest to investigate possible genetic differentiation between individuals collected during the late-winter/early-spring peak and those from the August peak. Because local bird communities differ between late winter/early spring—when most migratory birds have not yet returned from southern regions—and August—when they have not yet begun their southward migration—the bird species used by nymphs for blood meals during these two periods may differ. This temporal difference may influence the dispersal of *I. frontalis* and the pathogens potentially acquired during blood feeding, given that bird species differ in their reservoir competence for tick-borne pathogens.

### Ixodes frontalis *larval phenology*

Using our semi-quantitative approach, we confirmed the larval phenology of *I. frontalis* previously reported by Agoulon et al. (2019) across three climatic regions, including two newly studied ones. In our study, larvae were completely absent from March–April to July (or even until September at site D, which had the lowest abundance of *I. frontalis* across all stages) and occurred in large numbers from September–October through February–March. Agoulon et al. (2019) reported a similar pattern during a three-year survey, with larval activity peaking in autumn–winter and sometimes thousands of larvae found on a single 10 m² transect. In the study by Remesar et al. (2023), *I. frontalis* larvae were recorded in Spain (Mediterranean climate) between October and January, with a peak in November—further confirming that this larval phenological pattern occurs over a wide geographic range.

### Comparative development time from the larval to the nymphal stage in I. frontalis and I. ricinus

Although the nymphal peaks of both *I. frontalis* and *I. ricinus* occur at roughly the same time of year—for *I. ricinus*, between March and May at sites near Carquefou (close to sites A and B) or Sénart (near site C) (Hoch et al. 2024; Wongnak et al. 2022)—the larval peaks occur at different times: summer for *I. ricinus* and autumn–winter for *I. frontalis*. This suggests that the development time from the larval to the nymphal stage is relatively short for *I. frontalis*—approximately five months for the primary nymphal peak in March or around ten months for the secondary peak in August—and longer for *I. ricinus* (about nine months between the larval peak in summer and the subsequent spring nymphal peak). This difference in phenology between the two species may arise from variability in the developmental duration of engorged larvae or differences in diapause length—developmental diapause in engorged larvae or behavioral diapause in newly molted nymphs.

Laboratory monitoring of engorged larvae—either collected from wild birds or obtained through experimental feeding—under semi-natural conditions could provide data on the duration required for molting into nymphs. Experimental manipulation of day length could also help identify potential photoperiodic cues that trigger the onset or termination of diapause in these ticks.

## Conclusion

Following our initial discovery of large numbers of *I. frontalis* larvae during autumn–winter beneath bamboo groves at a first location (Agoulon et al. 2019) and the subsequent confirmation that this habitat is particularly suitable for the species across numerous localities (Plantard et al. 2021), we applied the accumulated knowledge to develop a new sampling protocol to further investigate the biology of this poorly known yet widely distributed tick. In this third study, the nymphal stage was also abundant in the same habitat, showing a pronounced peak in March. This finding paves the way for future investigations on *I. frontalis* that require substantial numbers of nymphs. Because most tick-borne pathogens are not vertically transmitted, larvae are pathogen-free; hence, nymphs are particularly suitable for assessing the prevalence of tick-borne pathogens potentially shared between *I. ricinus*—a species that commonly bites humans—and *I. frontalis*, which feeds exclusively on birds. Although our new protocol enabled the collection of a large number of *I. frontalis* nymphs within a limited sampling effort, adult *I. frontalis* remained scarce (11 females and 9 males during one year of monthly sampling). Although these numbers exceed those reported in previous studies—five adults over three years of monthly sampling in Agoulon et al. (2019) (1 in August, 3 in September, 1 in October) and one adult in one year of sampling in Remesar et al. (2023, with no timing provided)—the maxima in our study (7 in May or 5 in September at site A; 2 in October at site B) remain insufficient to confidently determine the timing of the adult peak. Therefore, additional investigations and the development of new approaches will be necessary to complete the picture of the full life cycle and phenology of this species.

## Acknowledgments

We thank the Parc floral de Paris, the cities of Nantes and Saint-Herblain and the private owner of the garden in the Prades-le-Lez for allowing us to collect ticks below bamboo. We are also grateful to the TMT (“Tiques et Maladies à Tiques” group) of the “Société Française d’Ecologie et d’Evolution” (SFE2; https://tmt.sfecologie.org/) for discussions and comments. The work of OP was funded by INRAE (Animal Health Division), CM and SM by ANSES, FS by CIRAD and AA by Oniris.

## Author contributions: CRediT

**Olivier Plantard :** Conceptualization, Formal analysis, Investigation, Methodology, Supervision, Visualization, Writing – original draft, review and editing, **Camille V. Migné :** Investigation, Writing – review and editing, **Sara Moutailler:** Investigation, Writing – review and editing, **Frédéric Stachurski:** Investigation, Writing – review and editing, **Albert Agoulon:** Conceptualization, Formal analysis, Investigation, Methodology, Supervision, Visualization, Writing – original draft, review and editing

